# Ribosomal RNA-specific antisense DNA and double-stranded DNA trigger rRNA biogenesis and insecticidal effect on insect pest *Coccus hesperidum*

**DOI:** 10.1101/2024.10.15.618468

**Authors:** Vol Oberemok, Nikita Gal’chinsky, Ilya Novikov, Alexandr Sharmagiy, Ekaterina Yatskova, Ekaterina Laikova, Yuri Plugatar

## Abstract

Invented in 2008, contact unmodified antisense DNA biotechnology (CUADb) is built on the use of short antisense DNA oligonucleotides (oligos) for insect pest control. Being a novel class of insecticides, oligonucleotide insecticides target pest rRNAs and/or pre-rRNAs and recently showed high insecticidal potential against sap-feeding insect pests, main vectors of plant DNA viruses and one of the most economically-damaging groups of herbivorous insects. In order to use all possible opportunities of CUADb, in this article insecticidal potential of short 11-mer antisense DNA oligos was investigated for *Coccus hesperidum* control in comparison with long 56-mer single-stranded and double-stranded DNA sequences and lower efficiency of the latter was found. At the end of the experiment, on the 9^th^ day, the highest mortality rate among antisense oligos was reached for Coccus-11 group (97.66 ± 4.04 %), while for long sequences the highest mortality rate was obtained for double-stranded DNA fragment in dsCoccus-56 group (77.09 ± 6.24 %). Also in this article architecture of DNA containment (DNAc) mechanism is described representing interesting and important for insect cell life interplay between rRNAs and different types of DNA oligos. During DNAc, Coccus-11 caused increased ribosome biogenesis and ATP production through metabolic switch in energy synthesis from carbohydrates to lipids but eventually caused ‘kinase disaster’ through downregulation of most kinases due to insufficient level of ATP produced. In the course of DNAc, hypercompensation of target rRNA is triggered by all highly and all somewhat complementary DNA oligos but more pronounced later degradation of target rRNA and significant insect pest mortality is seen only in the case of perfect complementarity of oligonucleotides to target rRNA. For both short and long oligonucleotide insecticides substantial decrease in rRNA concentration after rRNA hypercompensation in average by 3.75-4.25 fold, is explained by the work of DNA-guided rRNase, like RNase H1. We detected significant upregulation of RNase H1 after application of Coccus-11 in this study. On the contrary, for short and long random DNA oligos concentration of rRNA decreased in average by 2-3 fold after rRNA hypercompensation due to normal half-life of rRNAs in insect cells provided by ribonucleases. Fundamentally important, obtained results show completely new principle of regulation of rRNA expression in the cell via complementary interaction between rRNAs and unmodified antisense sequences of exogenous DNA. Practically important, this minimalist approach of using short antisense DNA dissolved in water is potent and eco-friendly innovation against sternorrhynchans and other pests, and reveals entirely new dimension to plant protection – DNA-programmable insect pest control.

## 1. Introduction

The Hemipterans, particularly sternorrhynchans, are recognized pests of many agricultural crops, being main vectors for plant DNA viruses and bacteria, and causing substantial yield losses [1,2]. Hemipteran pests are mainly controlled by neonicotinoids, leading to the evolution of resistance in field populations [3-6]. Also hemipterans can feed on plants and excrete honeydew contaminated with neonicotinoids at different concentrations that may cause lethal and sublethal effects on beneficial insects [7]. Emerging genetic resistance to chemical insecticides is constantly pushing for the search of new insecticides [8,9]. History of use of chemical insecticides indicates need for a major paradigm shift in the development of insecticides in order to obtain controlling agents with multi-decade utility and has been an important goal for many decades.

In 2008, an entirely new dimension of insect pest control – DNA-programmable plant protection – was discovered when unmodified DNA was shown to possess insecticidal effect [10]. Since then, the main breakthroughs in this direction were in finding the most convenient target genes (rRNA genes), in revealing mechanism of action (DNA containment) and in searching up insect pests (sternorrhynchans) with high susceptibility to this approach [11]. Fundamentally, for the first time it was found that short unmodified antisense DNA can both up-regulate and then down-regulate expression of rRNA genes controlling 80% of cell RNA. Up-regulation and further down-regulation of expression of rRNA can be essential for rDNA transcription master regulation during rRNA expression, regulation of rRNA expression during replication of DNA viruses, antiviral defense against DNA viruses [12].

In the last few years, contact unmodified antisense DNA biotechnology (CUADb) based on oligonucleotide insecticides (briefly, olinscides or DNA insecticides) has been established as a potent and selective ‘genetic zipper’ method (a target rRNA and/or pre-rRNA and an olinscide interlock and in the presence of DNA-guided rRNase resemble zipper mechanism performed by DNA–RNA duplex) [13]. CUADb-based ‘genetic zipper’ method is used for control of many hemipteran pests, particularly sternorrhynchans: soft scale insects, armored scale insects, whiteflies, psyllids, mealybugs, and aphids, as well as other groups of pests, thrips and spider mites, opening new horizons in green plant protection based on deoxyribonucleic acid. Recently, in Frontiers of Agronomy opinion article was published on successful list of pests targeted by oligonucleotide insecticides [14]. All 13 oligonucleotide pesticides represented in this article were selected at the first attempt according to the same algorithm and did work very efficiently; in average, the developed olinscides caused 80.04 ± 12.73% mortality for sternorrhynchans in 3–14 days after single or double treatment. Because of very efficient and easy algorithm, DNA-guided ‘genetic zipper’ method (CUADb) is a unique and very potent alternative to other approaches in plant protection based on duplexes of unmodified nucleic acids and RNA-guided nucleases: RNA interference and CRISPR/Cas9 [15].

While RNAi and CRISPR/Cas9 were not discovered on insect pests and initially had fundamental importance rather than practical one, CUADb was discovered on insect pests as practical tool and recently fundamental importance of this phenomenon for rRNA biogenesis was revealed [11,14,16-18]. To date, while RNAi and CRISPR/Cas9 are excellent tools for manipulations with unmodified nucleic acids in laboratory, they do not have easy algorithms for creation of end-products for plant protection; each separate case is special and usually is sorted out using trial and error method [19,20].

Ribosomal RNAs of insect pests serve as convenient targets for oligonucleotide insecticides, since rRNA makes up 80% of all RNA in the cell; rRNAs are ‘house-keeping’ non-coding RNAs that have important roles in the cytoplasm and beyond, including regulation of translation, signaling and metabolism [21]. Thousands of different mRNAs make up only 5% of all RNA and use of mature rRNA and pre-rRNA for targeting substantially increases signal-to-noise ratio, ca. 10^5^:1 (rRNA vs. random mRNA) [22]. Thus, it is almost impossible to get substantial insecticidal effect with an oligonucleotide insecticide targeting a random mRNA, still exclusions can be found when a particular gene is strongly activated, for example, IAP genes during baculovirus infection [23], or when a group of genes are targeted by one antisense oligonucleotide. Oligonucleotide insecticides act through DNA containment mechanism (DNAc) [17,18] which turned out to be highly significant and conserved in hemipterans. DNAc is a 2-step mechanism discovered on sternorrhynchans that takes place in nucleus [17], precisely nucleolus, where ribosome biogenesis takes place [24]. At first step of DNAc, target rRNA is ‘arrested’ and hypercompensation of target rRNA occurs (protein machinery and precise architecture of this step remains a riddle). At second step of DNAc, target rRNA degradation by DNA-guided rRNases, like RNase H1 takes place [18]. Formation of DNA olinscide-rRNA duplex resembles zipper mechanism. This duplex contains (or ‘zips’) normal target rRNA expression and leads to death of pests. Above-mentioned approach also was termed as ‘genetic zipper’ method [13].

In our opinion, partly, oligonucleotide insecticides are efficient against sternorrhynchans due to anatomic peculiarities of these pests: they contain numerous spiracular pores, preopercular pores, simple pores, etc. [25] that obviously enable deeper penetration of DNA oligos into the organism. Contact oligonucleotide insecticides, as the next-generation class of insecticides [26], are supposed to have fast biodegradability, selectivity in action, low carbon footprint, and avoid target-site resistance [18]. Oligonucleotide insecticides for sternorrhynchans represent minimalist approach, just short antisense DNA dissolved in water, and have minimal risks for the environment. They can resolve the problem of developing target-site resistance to insecticides in insect pests [27], if unique and highly conservative sequences of rRNA genes will be used. If insecticide resistance occurs, new and efficient olinscides can be easily re-created displacing target site to the left or to the right from the olinscide-resistance site of target mature rRNA and/or pre-rRNA [13,18]. Particularly important is necessity of complementarity of 3′-end nucleotide of olinscide to ensure maximum insecticidal effectiveness of oligonucleotide insecticides (3’-end rule of olinscides) [18]. Importantly, non-canonical base pairing, such as A:C (C:A) and G_olinscide_:U_rRNA_ [18,28,29] may occur between DNA olinscides and imperfect sites of rRNAs and should be taken into consideration during design of oligonucleotide insecticides [18]. Liquid-phase DNA synthesis based on phosphoramidite chemistry [30,31] makes it possible to significantly reduce the cost of this approach [32], providing 0.5 USD/ha for the ‘genetic zipper’ method in aphid control [15] and competing with current chemical insecticides. Therefore, the qualities of unmodified nucleic acids used as pesticides appear to be very promising [33,34].

Our research group proposed the contact use of short antisense DNA in plant protection in 2008 [10], rethought DNA-programmable insect pest control in 2019 [14], and continues to explore the facets of CUADb to find the most optimal formulations. In general, oligonucleotide insecticides are very effective on representatives of the order Hemiptera [35], and also moderately effective on representatives of the order Lepidoptera [23,36] and Coleoptera [37]. Already today, according to some estimations, ‘genetic zipper’ method is capable of successfully controlling 10-15% of the most serious insect pests of the world [15]. This approach is successfully developing to new groups of pests [11,38,39] and also helps to create species-specific and interspecific mixtures of oligonucleotide insecticides against mixed insect pest populations [18]. Addition of auxiliary substances (spreaders, adhesives, penetrators, and UV protectants) to formulation is possible in each separate case and their effect on efficiency and safety of the final formulation should be previously evaluated. Oligonucleotide pesticides are also compatible with the use of viral [23] and fungal [38] biopesticides showing higher mortality of the pests. It was found that contact delivery of unmodified antisense DNAs (CUADs) is much more efficient than oral delivery of unmodified antisense DNAs (ODUADs) because of active DNases present in digestive tract of insects [40].

Of note, recent results demonstrated remarkable specificity of oligonucleotide insecticides in action [36,41] and showed their safety for several non-target organisms: *Quercus robur, Malus domestica* [42], *Triticum aestivum* [43,44], *Manduca sexta, Agrotis ipsilon* [45], *Galleria mellonella* [36].

In this article, brown soft scale insect *Coccus hesperidum* L. (Hemiptera: Coccidae) was used as a model object. Brown soft scale is probably one of the most polyphagous insects, having been recorded from host plants in 345 genera belonging to 121 families. *C. hesperidum* may have originated from South Africa; it has spread through trade in infested plant material. Brown soft scale feeds on phloem sap, producing sugary honeydew waste that coats nearby plant surfaces and often gives rise to sooty mold growths. The main damage is indirect, when sooty molds coat the leaves, flowers and fruit; this blocks light and air from the leaves, impeding photosynthesis and respiration and modifying other physiological processes, resulting in loss of vigor and reduction in the size of fruit. *C. hesperidum* is not on any plant quarantine lists, because the species is so widespread already. Brown soft scale is a common multi-crop pest in greenhouses and is well-known as the main coccid pest of citrus with an extremely wide host range such as olive, avocado, cotton, mango, cocoa, ficus, hibiscus, oleander, palm, fern and orchid [46].

For this research, it was decided to use short (11-mer) and long (56-mer) 1) singlestranded and double-stranded DNA fragments with perfect complementarity and 2) somewhat complementary DNA fragments to target 28S rRNA of *C. hesperidum*. Evaluation of their efficiency as well as understanding of their action on insect cells is the main goal of the article.

## 2. Results

### 2.1. Mortality of C. hesperidum after treatment with single-stranded and double-stranded DNA sequences in the natural habitat

Insecticidal potential of 11-mer oligonucleotide insecticide Coccus-11 was evaluated on the viability of *C. hesperidum* larvae and caused significant 48.08 ± 6.56 % pest mortality (χ2 = 145.79, p < 0.001, N = 600, df = 1) on the 2^nd^ day compared to water-treated Control (Figure 2). Similar results, 66.33 ± 2.51 % pest mortality, was obtained on the 2^nd^ day for oligonucleotide insecticide Coccus_(−2)_-11 (χ2 = 249.64, p < 0.001, N = 600, df = 1) (Figure 1). Mortality progressively increased in Coccus-11 group up to the end of the experiment. By the 6^th^ and 9^th^ days of the experiment, pest mortality in Coccus-11 group reached 88.33 ± 13.42 % (χ2 = 401.61, p < 0.001, N = 600, df = 1) and 97.66 ± 4.04 % (χ2 = 463.62, p < 0.001, N = 600, df = 1), respectively. Similar trend was obtained for oligonucleotide insecticide Coccus_(−2)_-11. By the 6^th^ and 9^th^ days of the experiment, pest mortality reached 91.11 ± 4.58 % (χ2 = 430.49, p < 0.001, N = 600, df = 1) and 96.20 ± 2.14 % (χ2 = 445.19, p < 0.001, N = 600, df = 1), respectively. Random oligonucleotide A_2_C_3_G_3_T_3_-11 did not show any significant insecticidal effect compared to water-treated Control, while random oligonucleotide CG-11 did show weak insecticidal trend. On the 9^th^ day of the experiment, pest mortality rate in A_2_C_3_G_3_T_3_-11 and CG-11 groups were 10.33 ± 1.53 % (χ2 = 0.02, p > 0.892, N = 600, df = 1) and 22.67 ± 2.52 % (χ2 = 17.75, p < 0.001, N = 600, df = 1), respectively (Figure 1).

**Figure 1.**
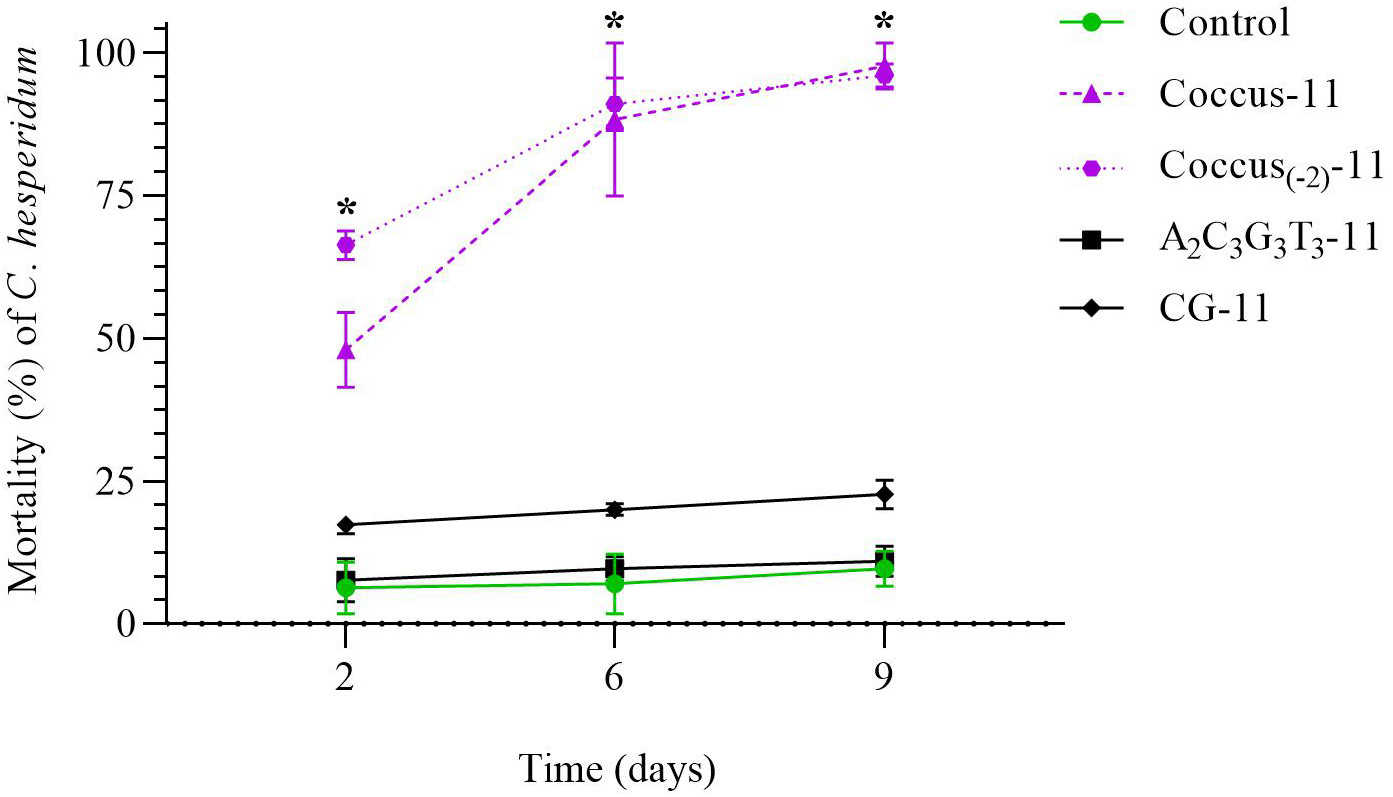
Dynamics of mortality of *C. hesperidum* after contact treatment with water, Coccus-11, Coccus_(−2_)-11, A_2_C_3_G_3_T_3_-11, and CG-11. The significance of differences in the groups of oligonucleotide insecticides (Coccus-11 and Coccus_(−2)-_11) compared to water-treated Control is indicated by * at p < 0.05.

**Figure 2.**
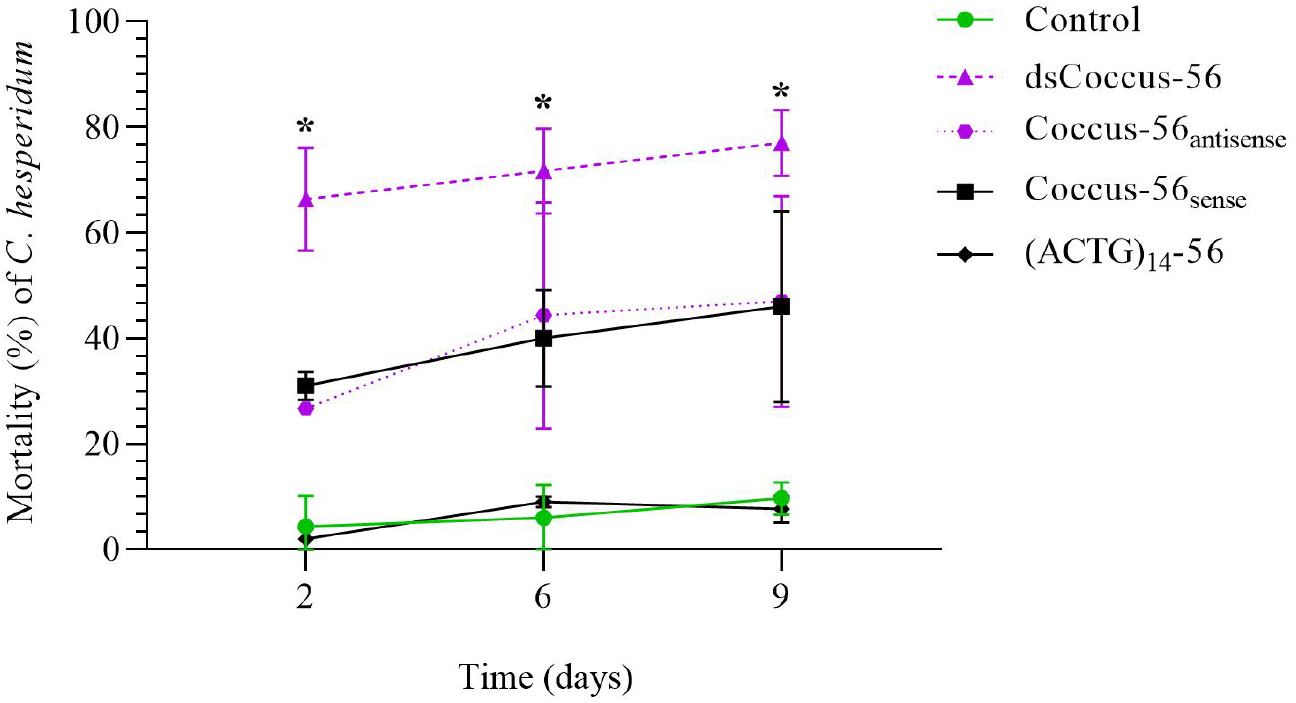
Dynamics of mortality of *C. hesperidum* after contact treatment with water, dsCoccus-56, Coccus-56_antisense_, Coccus-56_sense_, and (ACTG)_14_-56. The significance of differences in the groups of the experiment (dsCoccus-56, Coccus-56_antisense_, Coccus-56_sense_) compared to the water-treated Control is indicated by * at p < 0.05.

The same trend was found for long DNA sequences. In dsCoccus-56 group pest mortality increased significantly and comprised 66.33 ± 9.71 % on the 2^nd^ day after treatment (χ2 = 249.64, p < 0.001, N= 600, df = 1) compared to water-treated Control (Figure 2). In the groups treated with Coccus-56_antisense_, Coccus-56_sense_, (ACTG)_14_-56 pest mortality comprised 26.67 ± 1.15 % (χ2 = 71.51, p < 0.001, N = 600, df = 1), 31.14 ± 2.64% (χ2 = 55.43, p < 0.001, N = 600, df = 1), and 2.33 ± 1.52% (χ2 = 3.87, p > 0.05, N = 600, df = 1), respectively.

On the 6^th^ day after treatment significant increase in pest mortality compared to water-treated Control was found in all experimental groups except (ACGT)_14_-56 group (dsCoccus-56, χ2 = 249.11, p < 0.001, N = 600, df = 18; Coccus-56_antisense_, χ2 = 96.12, p< 0.001, N = 600, df = 1; Coccus-56_sense_, χ2 = 100.27, p < 0.001, N =600, df = 1; (ACGT)_14_-56, χ2 = 1.53, p > 0.215, N = 600, df = 1). On average, 7.33 ± 4.93 %, 71.68 ± 8.17%, 44.33 ± 21.38 %, 40.05 ± 9.2 %, and 9.26 ± 1.43 % of insects died on the 6^th^ day in groups of water-treated Control, dsCoccus-56, Coccus-56_antisense_, Coccus-56_sense_, and (ACGT)_14_-56, respectively (Figure 2).

On the 9^th^ day of the experiment, pest mortality was significantly higher in all experimental groups compared to water-treated Control (except (ACGT)_14_-56 group) and reached 9.68 ± 3.1 % in water-treated Control, 77.09 ± 6.24 % in dsCoccus-56 group (χ2 = 250.15, p < 0.001, N = 600, df= 1), 47.36 ± 19.97 % in Coccus-56_antisense_ group (χ2 = 96.78, p < 0.001, N = 600, df = 1), 46.42 ± 18.02 % in Coccus-56_sense_ group (χ2 = 101.13, p <0.001, N = 600, df = 1), and 7.68 ± 2.51 % in (ACGT)_14_-56 group (χ2 = 0.52, p > 0.469, N = 600, df = 1) (Figure 2).

Both short (Coccus-11, Coccus_(−2)-_11) and long (dsCoccus–56, Coccus-56_antisense_) oligonucleotide insecticides triggered significant pest mortality. In our opinion, moderate insecticidal potential of Coccus-56_sense_ is explained by its interference with normal interaction of native 28S rRNA and ribosomal proteins, in the same manner found for antibiotic binding sites [47]. Generally, short olinscides showed greater insecticidal potential in comparison with longer ones. Though olinscide dsCoccus-56 caused substantially higher pest mortality by 39.61 ± 0.91 % in comparison with Coccus-56_antisense_ (χ2 = 46.69, p < 0.001, N =600, df = 1) and Coccus-56_sense_ (χ2 = 43.46, p < 0.001, N =600, df = 1), it was in average by 21.11 ± 0.07 % lower than for short olinscides Coccus-11 (χ2 = 68.42, p < 0.001, N =600, df = 1) and Coccus_(−2)_-11 (χ2 = 56.43, p < 0.001, N =600, df = 1) at the end of the experiment.

The highest pest mortality was seen only for short and long olinscides and occurred between 2^nd^ and 6^th^ days. Of note, among all investigated short and long olinscides, dsCoccus-56 had the highest insecticidal effect (66.33 ± 9.71 % mortality) on *C. hesperidum* on the 2^nd^ day. On the 6^th^ day, short olinscides (Coccus-11 and Coccus_(−2)-_11) significantly increased their insecticidal effect on the pest and surpassed dsCoccus-56 in efficiency (88.33 ± 13.42 % and 91.11 ± 4.58 % vs. 71.68 ± 8.17 % mortality, respectively) (p<0.05). It seems advisable to use formulations of oligonucleotide insecticides against *C. hesperidum* containing both long double-stranded and short single-stranded DNA sequences. This will allow a rapid (in 1-2 days) insecticidal effect to be achieved with double-stranded olinscides, and the remaining pest population will be reduced with short olinscides in a few days after.

### 2.2. Target rRNA expression of C. hesperidum during DNAc

During DNA containment mechanism, hypercompensation of target rRNA was triggered by all DNA oligos (Figures 3-4, a; 6). Hypercompensation of 28S rRNA progressively increased from the 2^nd^ to the 6^th^ day (peak for all experimental groups, except Coccus-11 that had peak on the 2^nd^ day) and then decreased by the 9^th^ day. Generally, short oligos (specific and random) triggered lower level of rRNA hypercompensation than long oligos. As a trend, short oligonucleotide insecticides (Coccus-11 and Coccus_(−2)_-11) also triggered the lowest level of rRNA hypercompensation (Figure 3, b) while long olinscides (Coccus56_antisense_ and dsCoccus56) triggered the highest (Figure 4, b). Short and long random oligos took an intermediate position in relation to level of rRNA hypercompensation (Figures 3-4, c).

**Figure 3.**
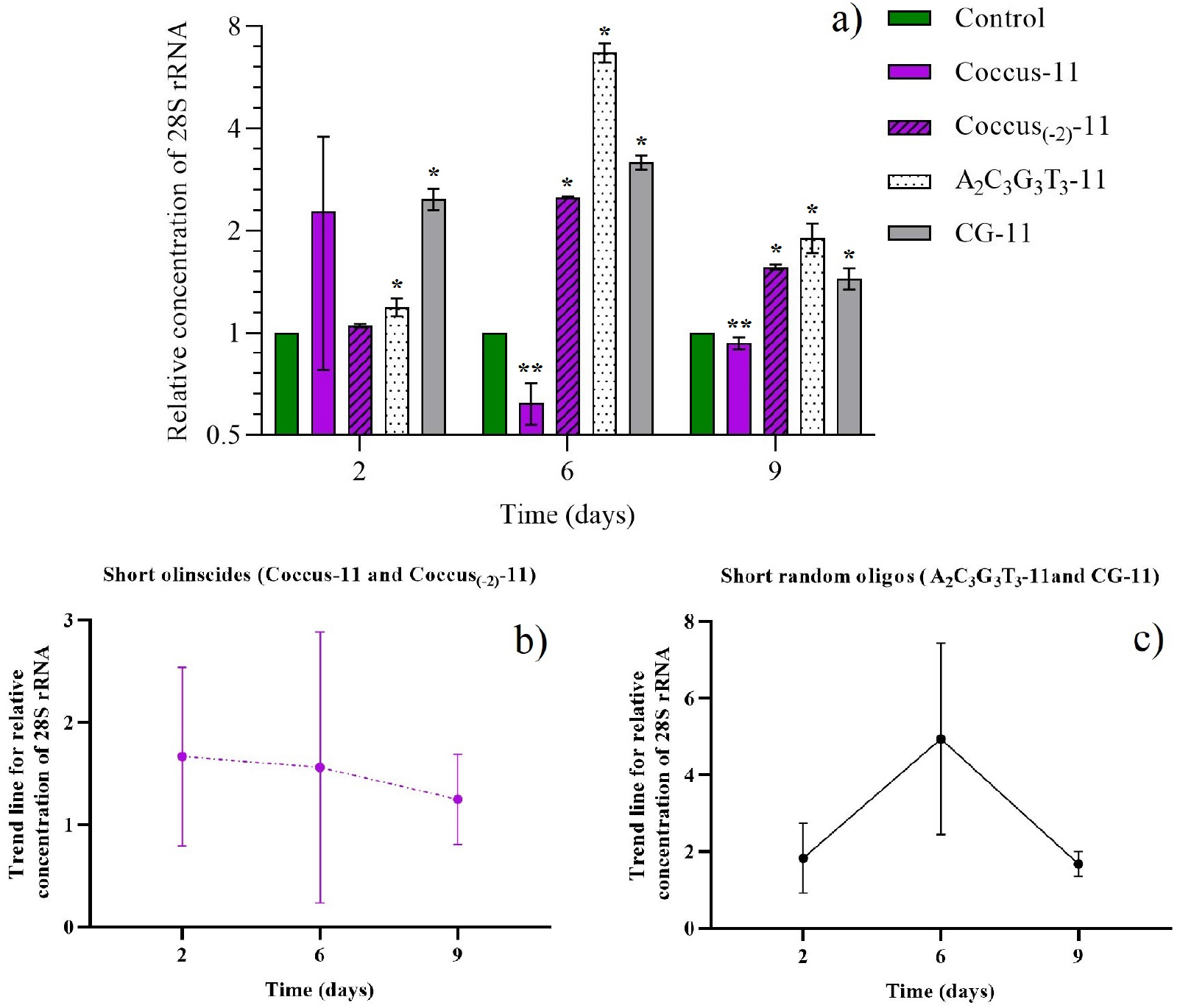
Dynamics of the relative concentration of 28S ribosomal RNA; a) for all short DNA oligos (short olinscides and short random oligos); b) for short olinscides (average concentration); c) short random oligos (average concentration); * is marked when concentration of target 28S rRNA is significantly higher compared to water-treated Control (p < 0.05); ** is marked when concentration of 28S rRNA is significantly lower compared to water-treated Control (p < 0.05); average relative concentration means that there are individuals in investigated bulk of insects with lower and higher concentrations of target rRNA in comparison with the average number.

**Figure 4.**
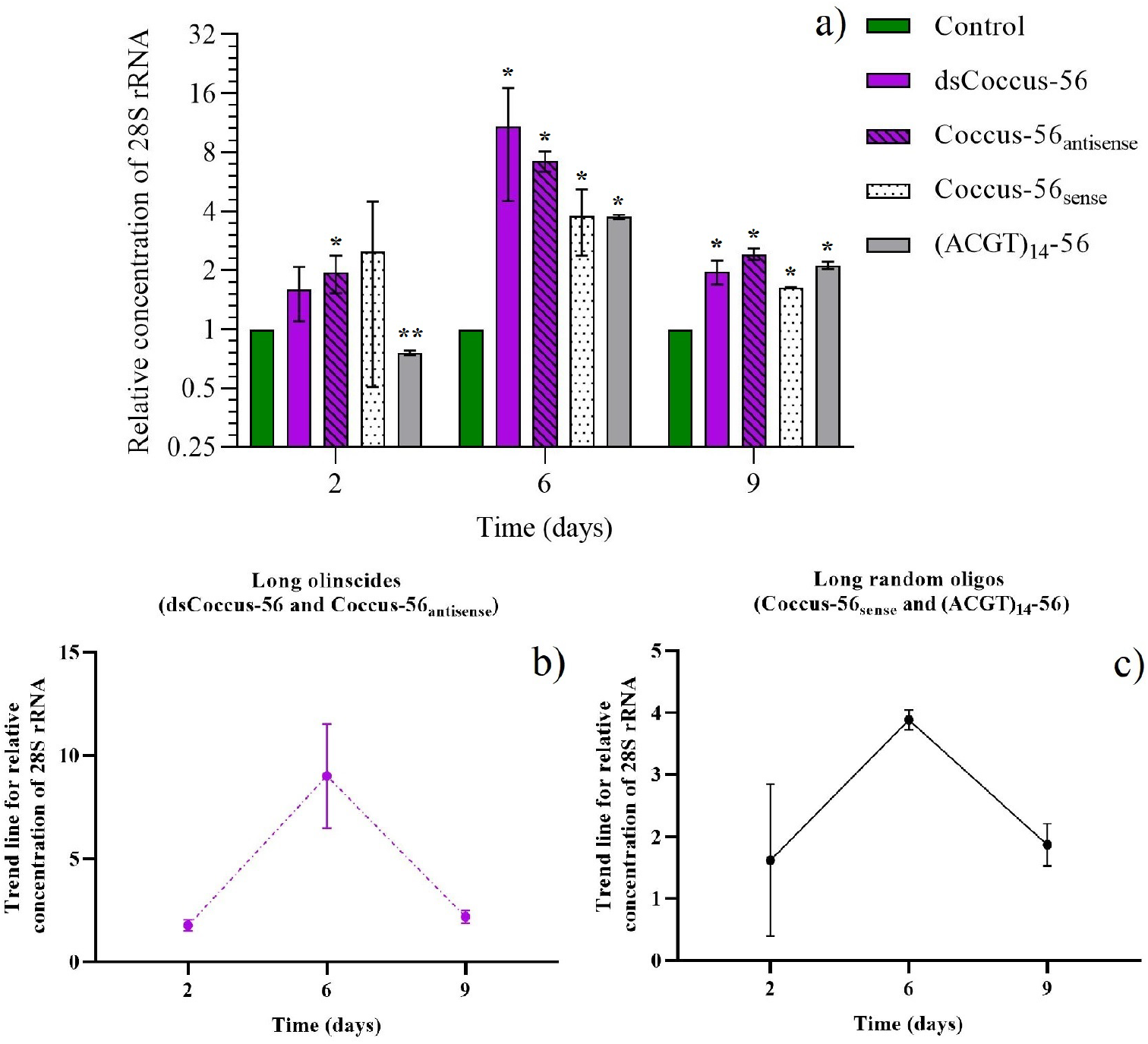
Dynamics of the relative concentration of 28S ribosomal RNA (average concentration) for long olinscides and long random oligos; a) for all long DNA oligos (long olinscides and long random oligos); b) for long olinscides (average concentration); c) long random oligos (average concentration); * is marked when concentration of target 28S rRNA is significantly higher compared to watertreated Control (p < 0.05); ** is marked when concentration of 28S rRNA is significantly lower compared to water-treated Control (p < 0.05); average relative concentration means that there are individuals in investigated bulk of insects with lower and higher concentrations of target rRNA in comparison with the average number.

For short olinscides in average 3.75 fold more decreased concentration of target rRNA compared to short random DNA oligos on 6^th^ day (peak of rRNA hypercompensation) was detected (Figure 3, b, c). In turn, for long olinscides 4.25 fold more substantial decrease in target rRNA concentration on 9^th^ day (after pronounced peak of rRNA hypercompensation on 6^th^ day) was detected (Figure 4, b, c). For both short and long olinscides substantial decrease in rRNA concentration after rRNA hypercompensation is explained by the work of DNA-guided rRNases, like RNase H1. For long random DNA oligos concentration of rRNA decreased by 2.04 fold and for short random DNA oligos concentration of rRNA decreased by 3.02 fold between 6^th^ and 9^th^ days, respectively, and this corresponds to normal half-life of rRNAs that lasts 3-5 days in cells and is degraded by ribonucleases [48].

Coccus-11 and Coccus_(−2)_-11 caused almost similar mortality but dynamics of rRNA expression in these two groups substantially differed. Only Coccus-11 down-regulated expression of target rRNA gene by 1.63 fold on the 6^th^ day. It can be explained by myriad binding partners of rRNA that limit its accessibility to antisense oligonucleotides and DNA-guided rRNase recruitment [49,50].

### 2.3. Histological studies

Histological studies were performed (Figure 5, a, b) to detect hypercompensation of rRNA in insect cells caused by DNA oligos. The 2^nd^ day of the experiment with short DNA oligos was chosen where substantial rRNA hypercompensation was detected in Coccus-11 group compared to water-treated Control. Upon histological examination with hematoxylin and eosin, in the Control group the apical parts of the insect cells were found to have outgrowths, in the recesses between which vesicles of the Golgi apparatus emerge with polysaccharides, forming the cuticle layer by layer. In insects of the control group, the network of such outgrowths is well defined, and the layering of the cuticle is also noticeable. From the basement of membrane side, the epithelium is washed by hemolymph (Figure 5, (b) – Control). Interestingly, in the A_2_C_3_G_3_T_3_-11 group, the color saturation of the cuticle was similar compared to water-treated Control group, and the thicknesses of cuticle layer was similar as well (Figure 5, (b) – A_2_C_3_G_3_T_3_-11). Insects from the A_2_C_3_G_3_T_3_-11 group show minor changes like slight thinning of the cuticle layer and loss of layering.

**Figure 5.**
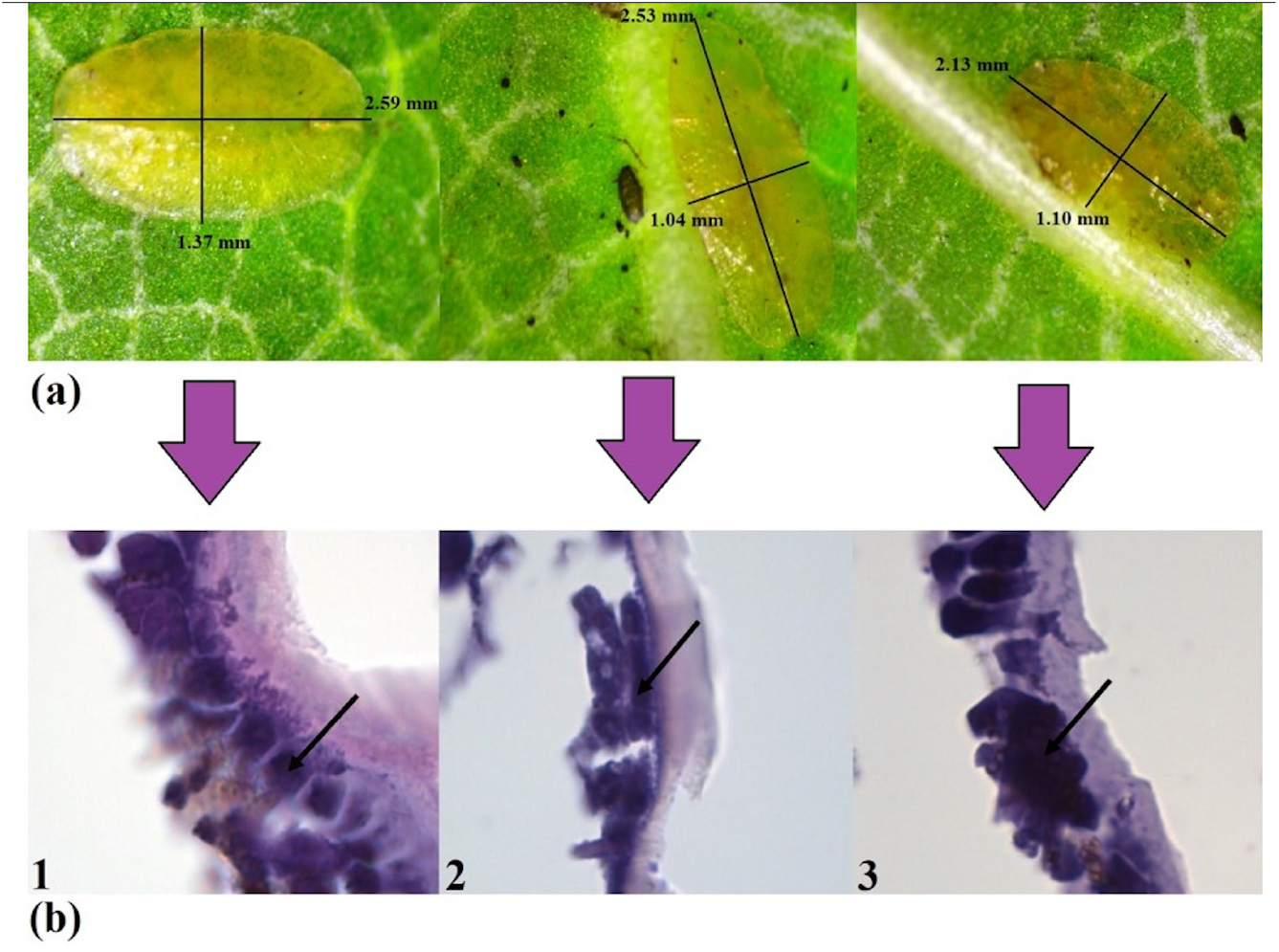
The effect of water (Control), random oligo (A_2_C_3_G_3_T_3_-11), and oligonucleotide insecticide (Coccus-11) on *C. hesperidum* larvae on the 2^nd^ day of the experiment; (a) insect morphology was investigated with light microscopy; (b) the integument of *C. hesperidum* is represented by a cylindrical single-layer epithelium from different groups of the experiment stained with hematoxylin and eosin; arrows show areas with the most intensive staining by hematoxylin (blue color) in each variant of the experiment.

In the Coccus-11 group, with a sufficiently high epithelium and preservation of the apical processes, the thickness of the cuticle is substantially smaller than in the control (Figure 5, (b)–Coccus-11). The cytoplasm of epithelial cells appears denser, with a large number of dark-stained hematoxylin granules. It is known that hematoxylin stains nucleic acids (DNA and RNA), the cell nucleus, ribosomes and RNA-rich areas of the cytoplasm [51]. Thus, such intensive staining of insect cells in the Coccus-11 group proves rRNA hypercompensation and increased level of rRNA biogenesis (found by RT-PCR; Figure 3, a) in comparison with water-treated Control and A_2_C_3_G_3_T_3_-11 groups on the 2^nd^ day after the treatment with DNA oligos (Figure 3, a).

### 2.4. Oligonucleotide insecticides are dominantly contact insecticides

During the experiment it was noticed that *C. hesperidum* occurred mainly on the abaxial side of *Pittosporum tobira* leaves Thunb. (Apiales: Pittosporaceae) in the investigated population. It was interesting to check whether olinscides have characteristics of systemic insecticides that move through plants [52]. To do this, adaxial side of the leaves without pest was treated with olinscide Coccus-11. As a control, abaxial side of *P. tobira* leaves containing pest also was treated. On the abaxial side of leaves, olinscide Coccus-11 caused substantial mortality of the pest (85.3 ± 8.9 %) compared to water-treated Control group (χ2 = 76.4, p< 0.001, N = 140, df = 1). Application of olinscide Coccus-11 on the adaxial side of leaves caused moderate mortality of the pest found on abaxial side of leaves and comprised 31.8 ± 7.1 % (χ2 = 11.6, p< 0.001, N = 133, df = 1) (Table 1).

**Table 1.** Mortality of *C. hesperidum* after application of water, random oligo A_2_C_3_G_3_T_3_-11, and olinscide Coccus-11 on the abaxial and adaxial sides of *P. tobira* leaves on the 4^th^ day.

| Adaxial side of the leaf |  |  | Abaxial side of the leaf |  |  |
| --- | --- | --- | --- | --- | --- |
| Control | A<sub>2</sub>C<sub>3</sub>G<sub>3</sub>T<sub>3</sub>-11 | Coccus-11 | Control | A<sub>2</sub>C<sub>3</sub>G<sub>3</sub>T<sub>3</sub>-11 | Coccus-11 |
| 7.2 ± 1.9 % | 7.9 ± 5.5 % | 31.8 ± 7.1 %* | 7.5 ± 2.4 % | 16.6 ± 6.8 % | 85.3 ± 8.9 %* |

\* – significance of the difference compared to water-treated Control, p<0.05

Random (somewhat complementary) oligonucleotide A_2_C_3_G_3_T_3_-11 did not show significant insecticidal effect compared to water-treated Control on either surface of leaves on the 4^th^ day. Thus, oligonucleotide insecticides based on unmodified antisense DNA do not possess pronounced characteristics of systemic insecticides and direct contact of oligonucleotide insecticides with insect integument is required for substantial insecticidal effect.

### 2.5. Differential gene expression analysis (DGE) after contact application of Coccus-11

On the 4^th^ day, DGE analysis of *C. hesperidum* in response to contact application of Coccus-11 was also performed in this study. This time period was chosen to cover the transition between the first (ribosome ‘arrest’ and target rRNA hypercompensation) and second (target rRNA degradation) steps of the DNA containment mechanism, when the maximum number of proteins are involved in the process (Figure 3, a). Particular attention was paid to the genes responsible for the biogenesis of ribosomal proteins and ribosomes in general, as well as genes involved in cellular energy production. It is known that ribosome biogenesis and maintenance of their functioning consumes 60% of the total cellular energy [53], so it was important to assess the balance in the functioning of these systems. In addition, attention was paid to the activity of RNase H1, as one of the known key enzymes involved in the degradation of target RNA using a complementary DNA sequence [54]. DGE analysis of the corresponding proteins would help to better understand the architecture of the DNAc mechanism against the background of an initial increase and subsequent decrease in the concentration of pest rRNA, as well as to better understand the causes of death of the insect pest (Table 2).

**Table 2.**
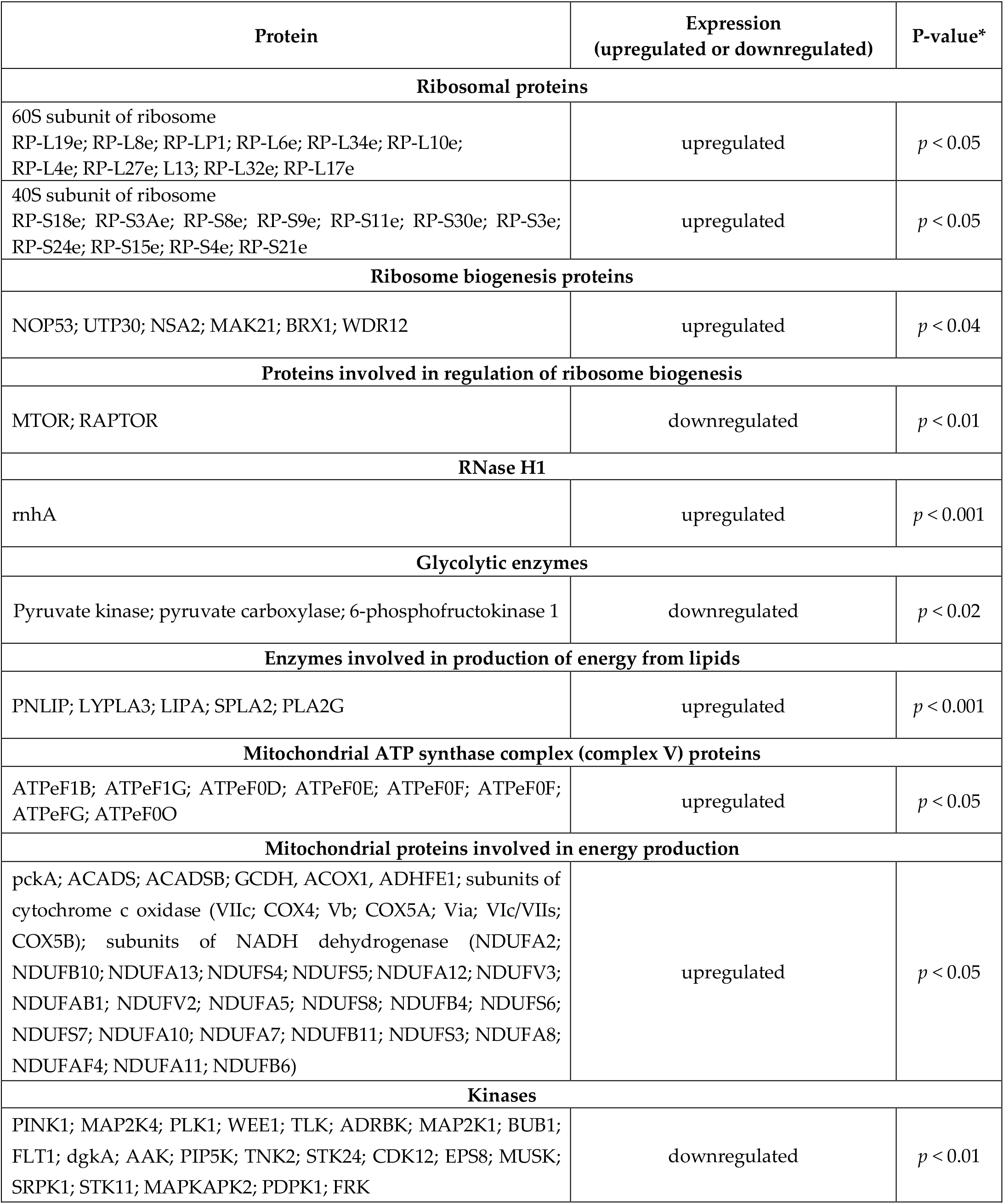
Differential gene expression (DGE) analysis of *C. hesperidum* after contact application of Coccus-11 vs. water-treated Control group on the 4t^h^ day; * difference in expression of each protein corresponds to the represented P-value.

On the 4^th^ day, it was found that, during DNAc, almost all ribosomal proteins of 40S and 60S ribosomal subunits were significantly upregulated promoting the formation of new ribosomes together with hypercompensated rRNA. Main ribosome biogenesis proteins [55] involved in the formation of ribosomal subunits (NOP53, UTP30, NSA2, MAK21, BRX1, WDR12) were upregulated as well. Also, during DNAc, most of ATP-dependent enzymes were downregulated (including mTOR, serine/threonine protein kinase which is together with Raptor is playing crucial role in ribosome biogenesis through mTORC1) [56] while proteins from mitochondrial ATP synthase complex and enzymes of mitochondrial complex crucial for maintaining cellular energy (phosphoenolpyruvate carboxykinase, cytochrome c oxidase, Acyl-CoA dehydrogenase, alcohol dehydrogenase, adenylate kinase, NADH dehydrogenase (ubiquinone), succinate-CoA ligase) [57,58] were significantly upregulated indicating deficiency of cellular energy caused by oligo-nucleotide insecticide Coccus-11. Moreover, during DNAc enzymes involved in production of energy from lipids [59] namely triacylglycerol lipase (PNLIP), lysophospholipase III (LYPLA3), lysosomal acid lipase/cholesteryl ester hydrolase (LIPA), secretory phospholipase A2 (SPLA2), PLA2G, were significantly upregulated. At the same time, crucial glycolytic enzymes [60,61] were downregulated (pyruvate kinase, aldolase, phosphofruc-tokinase-1) while none of glycolytic enzymes was upregulated, indicating a metabolic switch in energy synthesis from carbohydrates to lipids possessing higher energy yield per unit of mass compared to carbohydrates. One more important observation was that RNase H1 was also significantly upregulated (2.4 fold) during DNAc. RNase H1 functions independently of cell cycle and cleaves RNA-DNA hybrids, including those formed between DNA and rRNA [62].

## 3. Discussion

### 3.1 Sequences of DNA oligos highly complementary to rRNA (olinscides)

In our studies on sternorrhynchans, particularly *C. hesperidum*, it was found that DNA containment mechanism consists of 2 steps and represents fundamental and previously unknown interplay between different types of DNA oligos (olinscides and random oligos) and rRNAs. The two-step DNAc mechanism has been clearly demonstrated in previous studies with oligonucleotide insecticides on *C. hesperidum* [16], *Trioza alacris* [13], *M. sanborni* [63], as well as on *Aonidia lauri* and *Dynaspidiotus britannicus* [18]. In this study, the working mechanism of DNAc has been demonstrated not only for short 11-mer DNA sequences but also for long 56-mer DNA fragments. The obtained results allowed to create a general scheme of action of target and random DNA oligonucleotides on sternorrhynchans, particularly on *C. hesperidum* (Figure 6). Ribosomal RNA plays role of a potent sensor for penetrated DNA oligos and comprises 80% of the cell RNA performing important functions during protein biosynthesis, apoptosis, DNA damage repair, etc. Thus, regulation of synthesis of rRNA is of immense value for cell life [64,65]. Obviously, DNA containment mechanism can serve as a module in DNA repair, virus-host relationship, action of extracellular DNA, regulation of rRNA expression by cell, etc. and is evolutionary conserved regulatory process of insects. At the same time, it is clear that the fine details of the influence of exogenous DNA on insect cells will continue to be studied. DNAc is the tip of the iceberg of the processes that take place in response to DNA oligos, especially in light of the protein machinery involved in this mechanism.

**Figure 6.**
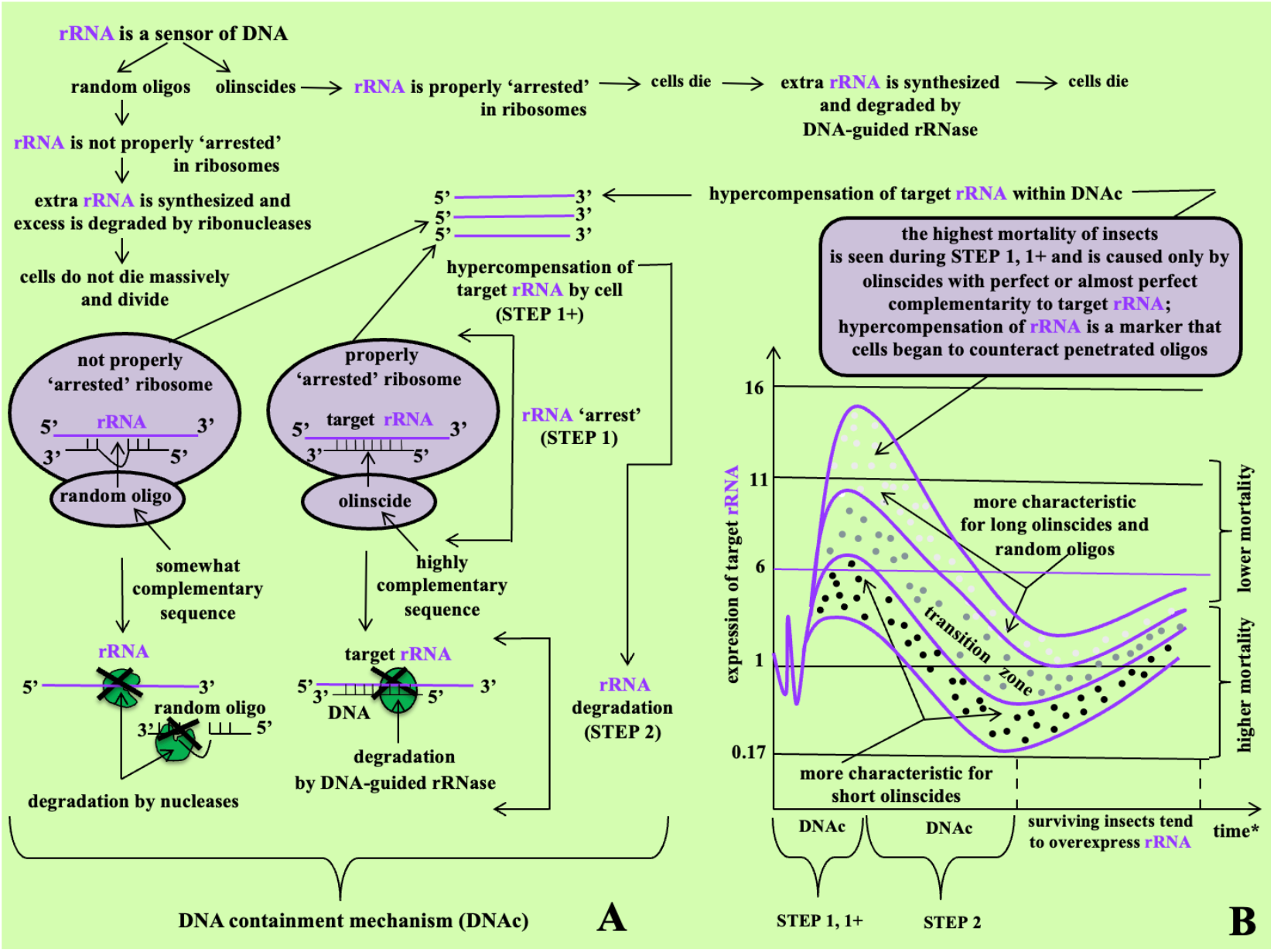
DNA containment mechanism (DNAc) of Sternorrhyncha representatives (A) and trends of target rRNA expression triggered by olinscides and random oligos during DNAc (B); * — time varies depending on distinct species and used oligos, usually 2 steps (full cycle) of DNAc occurs within 2 weeks; transition zone — can be detected less frequently for short olinscides, long olinscides and random oligos.

At first step of DNAc, an antisense DNA oligonucleotide (olinscide) complementarily interacts with target mature rRNA (in other words, blocks target rRNA) and/or 47S pre-rRNA, thus, interferes with normal functioning of mature ribosomes and processing of 90S pre-ribosomes; this process is accompanied with substantial insect pest mortality. Target rRNA hypercompensation by the DNA-dependent RNA polymerase (Pol I) in insect cells occurs in response to proper ‘arrest’ of target rRNA by antisense DNA oligonucleotide. Both ‘old’ blocked target rRNA in ribosomes (ribosomes apparently do not function properly in this situation) and ‘newly’ synthesized target rRNA and polycistronic rRNA transcripts (47S pre-rRNA) can be found by RT-PCR and, thus, target rRNA hypercompensation is detected. Precise architecture of this step remains a riddle. At second step of DNAc, DNA-guided rRNase, like RNase H1, cleaves target rRNA and substantial decrease in its concentration occurs; this process is also accompanied with substantial insect pest mortality. Both short and long olinscides, either constantly keep the target rRNA concentration low (Coccus-11 and Coccus_(−2)_-11 in comparison with short random DNA oligos), or they dramatically reduce it after the peak of rRNA hypercompensation (Coc-cus56_antisense_ and dsCoccus56 in comparison with long random DNA oligos), which in both cases is explained by the work of DNA-guided rRNase. Obviously, DNA-guided rRNase protects its unmodified DNA guide from degradation by DNases.

This study has made significant progress in deeper understanding of DNAc mechanism. The results indicate that in response to the use of the oligonucleotide insecticide Coccus-11, increased expression of ribosomal proteins promotes creation of new ribosomes together with hypercompensated rRNA. At the same time, increased synthesis of ATP by mitochondria occurs mainly due to lipid degradation but eventually leads to ‘kinase disaster’ through downregulation of most kinases due to insufficient level of ATP produced. Obviously, a lack of ATP and decreased activity of kinases leads to exhaustion and death of the insect pest. Kinases are involved in approximately 50% of all cellular reactions and play a crucial role in cellular signaling and regulation by transferring phosphate groups from ATP to other molecules (phosphorylation). This phosphorylation can modify the activity, localization, and interactions of the target protein, thus affecting a wide range of cellular processes. [66,67] The obtained data correspond to the DNAc mechanism, core process of which is initial hypercompensation of rRNA to create new ribosomes, followed by rRNA degradation recruiting DNA-guided rRNases, like RNase H1 which was significantly upregulated during DNAc in this study. Further studies will show the response of a wider range of enzyme systems to oligonucleotide insecticides during DNAc, and will also show specific changes in DNAc when targeting, for example, mitochondrial rRNAs or ITS regions of pre-rRNA.

As a working hypothesis, DNA is not only template for rRNA synthesis, it is also direct regulator of rRNA expression. DNA containment mechanism (STEP 1+, hypercompensation of rRNA) can play crucial role in regulation of rRNA expression by complementary cell DNA (direct rDNA transcription master regulation) and viral DNA (rRNA switchboard mechanism) [11] as well as can be recruited in innate immunity system [18] against ssDNA viruses for which hemipteran insects serve as major vectors [2,68] and also against DNA viruses that normally infect them [50]. Insect nucleases can cleave specific DNA sequences of the invader producing ssDNA oligonucleotides and subsequently initiate degradation of target viral RNAs [18] (Figure 7). Obtained here results with unmodified DNAs (single-stranded and double-stranded) is a profound model for the described process and entirely new dimension to regulation of rRNA genes controlling 80% of cell RNA.

**Figure 7.**
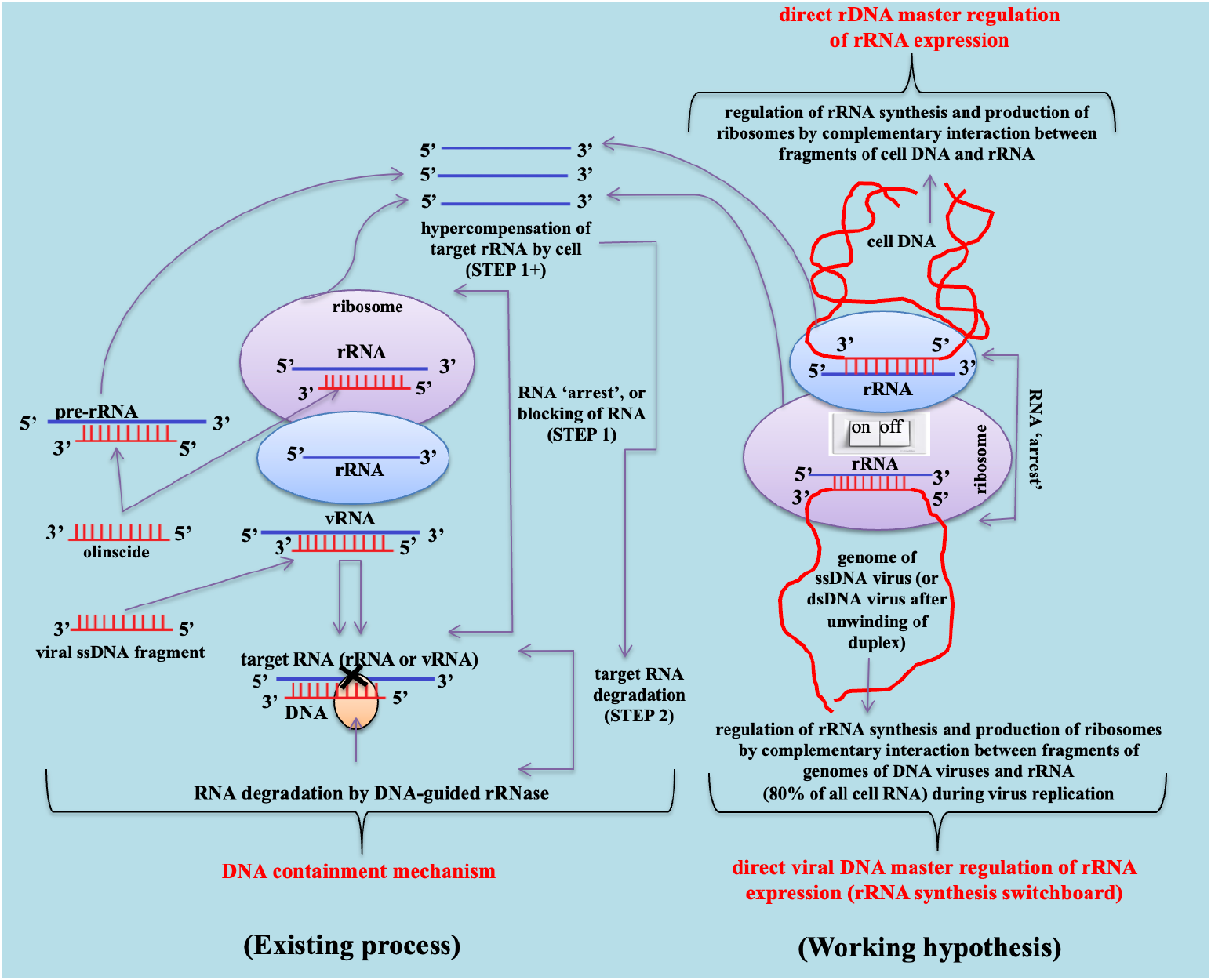
Mode of action of oligonucleotide insecticides, viral single-stranded DNA fragments, regulation of rRNA synthesis via direct rDNA transcription master regulation and direct viral DNA master regulation of rRNA expression based on DNA containment mechanism.

### 3.2. Sequences of DNA oligos somewhat complementary to rRNA (random oligos)

At first step of DNAc, a random (somewhat complementary) DNA sequence in imperfect manner interacts and not properly blocks target mature rRNA and/or 47S pre-rRNA, as a result, weakly interferes with normal functioning of mature ribosomes and processing of 90S pre-ribosomes.

Obviously, in this case, target rRNA hypercompensation by the DNA-dependent RNA polymerase is the reaction on DNA oligo as possible damage of cell DNA. Insect cells mobilize their strengths through elevation of protein biosynthesis (for example, it is known that pre-ribosomal RNA reorganizes DNA damage repair factors in nucleus during meiotic prophase and DNA damage response) [69]. Both weakly arrested ‘old’ RNA in ribosomes and ‘newly’ synthesized target rRNA and polycistronic rRNA transcripts (47S pre-rRNA) can be found by RT-PCR and, thus, was detect target rRNA hypercompensation. At second step of DNAc, ribonucleases cleave excess of rRNAs and deoxyribonucleases cleave penetrated DNA oligos. rRNAs have 3-5 days long cellular half-lives [47] and substantial 2-3 fold decrease in concentration of rRNA between 6^th^ and 9^th^ days was detected without significant pest mortality for groups with random DNA oligos. Obviously, DNA oligos without perfect complementarity to target rRNAs can’t recruit DNA-guided rRNase efficiently, eventually insect cells normalize their life through degradation of random DNA oligos. Generally, random oligos trigger cell proliferation and increase in cell volume, cells do not die massively, and substantial mortality of insects does not occur [70].

Thus, short and long single-stranded DNA oligonucleotides having weak interaction with target rRNA also trigger its hypercompensation. Non-specific interactions become inevitable with the increase of participating biomolecules and cell requires very clear signal – perfect or almost perfect complementarity of a DNA oligo to target rRNA – of whether to die or not. All other weak signals which are caused by random DNA oligos are subject to homeostasis. Subsequently, significant pest mortality is caused only by olinscides with perfect or almost perfect complementarity to target rRNA [18]. Obtained results open up vast opportunities for potent and selective insect pest control as well as reveals completely new principle of biogenesis of the most abundant RNA in the cell – rRNA.

### 3.3. Horizons of fundamental understanding and practical application of obtained results

In our opinion, closely related representatives of superorder Hemipteroidea (Paraneoptera) as well as groups of insect pests that can be infected by DNA viruses and actively involved in the transmission of plant DNA viruses will be the most susceptible to oligonucleotide insecticides. This is because they apparently have a natural DNAc defense mechanism against DNA viruses that helps them track and degrade viral mRNAs [18]. Vice versa, DNA viruses also can use DNAc mechanism (hypercompensation of rRNA) in order to increase the number of rRNAs for extra ribosomes in the cell necessary for their replication. The rapidly accumulating amount of genomic information indicates that a noticeable number of viral genomes contain sequences related to ribosomal RNA [71]. Also complementary sequences of cell DNA could regulate rRNA expression through formation of DNA-rRNA duplexes and initiation of DNAc mechanism. It is important to note that cell DNA and viral DNA do not take any risk to be degraded when they recruit DNA-guided rRNase, while rRNA can be either up-regulated or down-regulated by complementary cell DNA and viral DNA as direct master regulator of rRNA biogenesis. Our discovery made with the help of unmodified DNAs reveals previously unknown dimension to gene regulation of rRNA genes controlling four fifths of all RNA in the cell.

To date, it is not clear why long olinscides trigger more pronounced rRNA hypercompensation than long random DNA oligos, before substantial decrease of target rRNA by DNA-guided rRNase occurs. Precise architecture of rRNA hypercompensation remains a riddle. It has been hypothesized that special proteins (rRNA sensing proteins) may control the proper functioning of rRNA by interacting with it. As previously mentioned, myriad binding partners of rRNA exist [49,50] and some of them can perform this function. When highly complementary DNA-rRNA duplex is formed, these rRNA sensing proteins could break off and transmit a signal to start transcription on rDNA. The more proteins break off and transmit the signal, the stronger rRNA hypercompensation is. Based on the experimentally found proportion in the rRNA hypercompensation peaks between 56 nucleotides long olinscides and water-treated Control on the 6^th^ day (ca. 9:1) (Figures 4,5), each such specific protein may interact with 6-7 nucleotides in rRNA. Concept of rRNA sensing proteins could explain why somewhat complementary DNA oligos trigger rRNA hypercompensation. Even complementarity of 6-7 nucleotides in a row will be enough to trigger rRNA hypercompensation. All tested here DNA oligos have this threshold level of complementarity to *C. hesperidum* 28S rRNA in GenBank for rRNA of *C. hesperidum* (found by BLAST program). rRNA looks like a sensor for highly and somewhat complementary sequences of DNA oligos and rRNA hypercompensation is one of the first cell reactions on them. Apparently, that is why Coccus56_sense_ does not trigger pronounced rRNA hypercompensation but causes significant pest mortality. It does both somewhat complementary interaction with rRNA and proper interaction with rRNA sensing proteins, thus, it interferes with normal interaction of native 28S rRNA and ribosomal proteins. Interestingly, short olinscides caused 5.76 fold lower hypercompensation of target 28S rRNA in comparison with long olinscides. Short olinscides are 5.1 fold shorter than long olinscides and obtained data corresponds to the hypothesis of existence of RNA sensing proteins. The longer is olinscide, the higher hypercompensation of target rRNA occurs.

Of note, long olinscides can serve as models (or prototypes) that simulate interplay between host rRNAs and DNA viruses that normally infect hemipterans [72]. As a trend, antisense sequences Coccus56_antisense_ and dsCoccus56 triggered 2.36 fold stronger hypercompensation of 28S rRNA on the 6^th^ day (peak of rRNA hypercompensation for all experimental groups, except Coccus-11) than Coccus56_sense_ and (ACGT)_14_-56. This phenomenon could be used by DNA viruses. It is obvious that DNA viruses can also take advantage of hypercompensation of rRNAs observed during the DNA containment mechanism in order to increase the number of rRNAs for extra ribosomes necessary for their replication [73]. Bioinformatics studies indicate that hijacking of host genes by viruses is common [71,74]. Indeed, during coevolution, DNA viruses could not miss the opportunity to hijack rRNA genes via lateral transfer of the nucleic acid from the host organism [75] as key players in metabolism, since rRNA makes up four fifths of all RNA in a cell [76], and more than 60% of all energy is spent on the production and maintenance of ribosomes [53]. Most DNA viruses replicate in nucleus and most viruses interact with the nucleolus that plays a major role in virus life and assembly of ribosomes [77]. The canonical function of nucleoli is ribosome biogenesis, where they support rRNA synthesis, processing, and assembly into pre-ribosomal subunits [24]. It is not difficult to imagine that a complementary fragment of DNA of virus genome can regulate host rRNA synthesis according to the ON/OFF principle, using this trigger to influence the cell (to turn on this mechanism and turn it off when required). This mechanism, deemed rRNA synthesis switchboard, can help virus initiate rRNA synthesis and the formation of new ribosomes with minimal energy consumption [17], using the complementary interaction of specific sequences of viral DNA and host rRNA. Fundamentally very important, found here potent effect of antisense unmodified DNA on rRNA shows potential of cell DNA in regulation rRNA expression. Thus, cell DNA may serve not only as the matrix for rRNA synthesis but as regulatory molecule of its expression. These findings significantly change our understanding of regulation of rRNA biogenesis

Practically, the introduction of CUADb-based ‘genetic zipper’ method into plant protection techniques will help create a flexible system for adaptation of oligonucleotide insecticides to the constantly changing genetics of pests during microevolution. Fundamental goal of short oligonucleotide insecticides – healthy crops without losing yield – helps to easily adapt to microevolution of pests and, to our latest projections, affordability of the approach makes it very perspective to be implemented in agriculture on a large scale. Chemical insecticides remain a cornerstone of insect pest management [78-82]. There are several key factors that drive the development of new classes of insecticides and the most important of these is economic cost of insect pest damage to agriculture and insecticide resistance which has dramatically and relentlessly increased since the mid-20th century [8,78,83-85]. The general mechanism underlying insecticide resistance is natural selection, which leads to an increase in frequency of specific alleles formed as a result of random mutations in insect pest population [86-88]. Antisense technologies (RNAi, CUADb, and CRISPR/Cas) are able to counteract insecticide resistance by targeting conserved genes or conserved gene regions and by facilitating the rapid development of effective pest control agents when resistance to existing insecticides emerges. While CUADb [14] and RNAi [89] demonstrate promising potential as the next-generation bioinsecticides, due to their fast biodegradability, selectivity, and low carbon footprint [11,90-99], CRISPR/Cas is primarily used to genetically attenuate insect pest populations through genetic engineering [100-109]. Nevertheless, these innovative antisense technologies and their combinations, offer an expansive repertoire for controlling insect pests, and the central challenge lies in selecting the optimal pest management strategy for each specific case.

## 4. Materials and Methods

### 4.1. Origin of C. hesperidum L

*C. hesperidum* larvae were identified on *P. tobira* in the Nikita Botanical Garden (Yalta, Republic of Crimea) (situated at 44°30′41.9′′N latitude and 34°13′57.3′′E longitude) and used them for the experiment. The treatment was carried out on *P. tobira* plants using a hand-held sprayer with a solution of oligonucleotides in water (100 mg/L). 1 mg DNA in 10 ml of solution per m^2^ of foliage containing the pest was used. Oligonucleotide insecticides were applied directly to 1^st^ and 2^nd^ stage larvae (accounting for about 80 % of all insect individuals), as well as *C. hesperidum* nymphs. Oligonucleotide insecticides were applied directly on foliage containing *C. hesperidum* changing the angle of attack of handheld sprayer so that the olinscide gets on the entire leaf surface of trees with insect pests located on them. During 8 replicates of the experiment, around 10400 larvae were treated in three independent experiments, and their survival rates were calculated statistically. Insects were counted for each replicate of each variant of the experiment on 20 leaves of *P. tobira*. Mortality was calculated on the 2^nd^, 6^th^, and 9^th^ days; the number of dead individuals was divided by the total number of individuals on the leaf and multiplied by 100%.

### 4.2. Sequences and applied short (Coccus-11; Coccus_(−2)_-11) and long (dsCoccus-56; Coccus-56_antisense_) olinscides

GenBank database was used (*C. hesperidum* isolate S6A395 large subunit 28S ribosomal RNA gene, partial sequence; https://www.ncbi.nlm.nih.gov/nuccore/MT317022.1, accessed on 27 March 2024) to design short (Coccus-11, 5′–CCA–TCT–TTC–GG–3′; Coccus_(−2)_-11, 5′–CAC–CAT–CTT–TC–3′, it is situated 2 nt to the right from Coccus-11 antisense sequence) and long (Coccus-56_antisense_, 5′–CCA–TCT–TTC–GGG–TAC–CAG–CGT–GCA– CGC–TGT–AGG-TGC–GCC–CCA–GTT–CGT–CGA–CGG–TC–3′ and its corresponding double-stranded copy, dsCoccus-56) sequences as contact oligonucleotide insecticides. DNA sequencing was also performed to check if oligonucleotide insecticides complementarily match with 28S rRNA sequences in investigated population of *C. hesperidum*. It was checked after DNA sequencing and it matched (data not shown). Olinscides were dissolved in nuclease-free water (100 ng/µL) and applied using a hand sprayer to *P. tobira* leaves (mg of olinscides per m^2^ of leaves), 10 mL of the solution was used per m^2^. As a control, a water-treated group was used.

### 4.3. Sequences and applied short (A_2_C_3_G_3_T_3_-11; CG-11) and long (Coccus-56_sense_; (ACGT)_14_-56) random oligos

As a control, short (A_2_C_3_G_3_T_3_-11, 5′–AAC–CCG–GGT–TT–3′; CG-11, 5′–CGC–CGC– CGC–CG–3′) and long (Coccus-56_sense,_ 5′–GAC-CGT-CGA-CGA-ACT-GGG-GCG-CAC-CTA-CAG-CGT-GCA-CGC-TGG-TAC-CCG-AAA-GAT-GG–3’; (ACGT)_14_-56, 5′–ACG-TAC-GTA-CGT-ACG-TAC-GTA-CGT-ACG-TAC-GTA-CGT-ACG-TAC-GTA-CGT-ACG-TAC-GT–3′) sequences were used. They were also dissolved in nuclease-free water (100 ng/µL) and applied using a hand sprayer to *P. tobira* leaves (mg of olinscides per m^2^ of leaves), 10 mL of solution was used per m^2^. As a control, a water-treated group was used.

### 4.4. Synthesis of oligonucleotides

All oligonucleotide sequences (without chemical modification) were synthesized on automatic DNA synthesizer ASM-800ET (BIOSSET, Novosibirsk, Russia) using standard phosphoramidite chemistry on the universal solid support UnyLinker 500 Å (ChemGenes, Wilmington, USA). Cleavage and deprotection were carried out at 55 ºC overnight using a concentrated solution of ammonia. The solution was then filtered and evaporated on a vacuum rotary evaporator (Heidolph Instruments GmbH & Co. KG, Schwabach, Germany). The resulting solid substance was dissolved in deionized water (Merck Millipore, Molsheim, France) to the required concentration, which was determined by measuring it on a spectrophotometer NanoDrop Lite (Thermo Fisher Scientific, Waltham, USA) [13].

### 4.5. Evaluation of 28S rRNA expression of C. hesperidum

*C. hesperidum* larvae were ground using a pestle in 1.5 mL tube to prepare them for RNA extraction by ExtractRNA kit (Evrogen, Moscow, Russia) according to the manufacturer’s instructions. Three independent extractions were carried out to produce the replicates for each treatment. For each extraction, 10 larvae were used from each group (Control, Coccus-11, Coccus_(−2)_-11, A_2_C_3_G_3_T_3_-11, CG-11, dsCoccus-56, Coccus-56_antisense_, Coccus-56_sense_, (ATCG)_14_-56. The quality and concentration of the extracted total RNA were assessed using a NanoDrop spectrophotometer (Thermo Scientific, Waltham, MA, USA). 1.5% agarose gel was used to run electrophoresis in TBE (Tris-borate-EDTA) buffer (10 V/cm) for 30 min, and 5 µL of the eluted RNA volume was loaded for each group [110]. Among all experimental groups, the quantity, intensity, and pattern of the RNA bands were equal.

For reverse transcription, the total RNA (50 ng) was annealed with Coccus-R primer (5′–ACG–TCA–GAA–TCG–CTG–C–3′) and analyzed using FastStart Essential DNA Green Master kit (Roche, Basel, Switzerland) according to the manufacturer’s instructions. The reaction was conducted at 40 °C for 60 min in a LightCycler® 96 (Roche, Basel, Switzerland). cDNA (2 µl) was added to the mixture with FastStart SYBR Green MasterMix kit (Roche, Basel, Switzerland). Primers, forward 5′–ACC–GTC–GAC–GAA–CTG–G–3′ and reverse 5′–ACG–TCA–GAA–TCG–CTG–C–3′, were used for quantitative real-time PCR studies to calculate the concentration of *C. hesperidum* 28S rRNA. The qPCRmix-HS SYBR (Evrogen, Moscow, Russia) master mix was used according to the manufacturer’s instructions. The following procedure of 10 min initial denaturation at 95 °C, followed by 30 cycles with 10 s denaturation at 95 °C, 15 s annealing at 62 °C, and 14 s elongation at 72 °C was used for amplification on a LightCycler® 96 instrument (Roche, Basel, Switzerland) [111]. PCR was repeated in triplicate for each condition. Finally, to estimate the specificity of amplification and the presence of additional products, all the PCR products were melted.

### 4.6. Histochemical Assay

Dehydration and paraffin impregnation were carried out in a Logos microwave histoprocessor (Milestone, Sorisole, Italy). Tissue blocks were sectioned into slices 4 µm thick and stained with hematoxylin and eosin. The following reagents for fixation, dehydration, wiring, and staining were used: formalin 10%, isopropyl alcohol, paraffin, o-xylene, and a set of dyes of hematoxylin and eosin (Biovitrum, St. Petersburg, Russia).

### 4.7. Differential gene expression (DGE) analysis

The quality and quantity of the isolated total RNA were checked using a BioAnalyser (Agilent Technologies, Inc., Santa Clara, USA) and an RNA 6000 Nano Kit (Agilent Technologies, Inc., USA). Total RNA was then used to obtain the polyA fraction using Dynabeads® mRNA Purification Kit (Thermo Fisher Scientific, Waltham, USA) according to the kit instructions. Libraries for massively parallel sequencing were then prepared from the polyA RNA using the NEBNext® Ultra™ II RNA Library Prep Kit (NEB) (New England Biolabs, Inc., Beverly, USA) according to the kit instructions. Library concentrations were determined using the Qubit dsDNA HS Assay Kit (Thermo Fisher Scientific, USA) on a Qubit 2.0 fluorometer (Thermo Fisher Scientific, USA). Library fragment length distribution was performed using the Agilent High Sensitivity DNA Kit (Agilent Technologies, Inc., USA). Sequencing was performed on a HiSeq1500 (Illumina, Inc., San Diego, USA) with the generation of at least 10 million short reads of 50 nucleotides in length. Reads were aligned to the genome using the STAR program and the calculation of differentially expressed genes was performed using the DESeq2.0 package (Bioconductor, Seattle, USA). DGE analysis was performed in two replicates for Coccus-11 and water-treated Control group with 100 insect individuals per each replicate in each group.

### 4.8. Statistical Analyses

The standard error of the mean (SE) was determined and analyzed using the Student’s t-test to evaluate the significance of the difference in 28S rRNA concentration between control and experimental groups on the 2^nd^, 6^th^, and 9^th^ days. The non-parametric Pearson’s chi-squared test (χ2) with Yates’s correction was performed to evaluate the significance of the difference in mortality between control and experimental groups on the 2^nd^, 6^th^, and 9^th^ days. All above-mentioned calculations were preformed using Prism 9 software (GraphPad Software Inc., Boston, USA).

## 5. Conclusions

Obtained results with antisense and double-stranded DNA fragments helped to create general architecture of interplay between rRNA and exogenous DNA based on DNAc mechanism, important for both practical application (DNA-programmable insect pest control) and fundamental understanding of functioning of the cell (regulation of rRNA synthesis). At first step of DNAc, target rRNA is ‘arrested’ and hypercompensation of target rRNA occurs. At second step of DNAc, target rRNA degradation by DNA-guided rRNase takes place. Tested here unmodified DNAs serve as a simple and robust models of investigation of complementary interactions between exogenous DNA (including viral DNA) and rRNAs unraveling new facets of crucial and previously unknown roles played by nucleic acids.

Insecticidal potential of short 11-mer antisense DNA oligos was investigated for *C. hesperidum* control in comparison with long 56-mer single-stranded and double-stranded DNA sequences and lower efficiency of the latter was found. Shorter sequences result in higher yields during DNA synthesis using phosphoramidite method, thus providing a greater mass of product and decreases production costs. Oligonucleotide insecticides primarily exhibit characteristics of contact insecticides rather than systemic ones. Despite not being systemic, oligonucleotide insecticides are effective by targeting specific rRNA sequences essential for insect survival, leading to their death. The obtained data show that the insect is not able to compensate for the effect of oligonucleotide insecticide due to increased protein biosynthesis against the background of ATP depletion and ‘kinase disaster’ in the pest cell during the DNA containment mechanism and eventually dies.

We are on the cusp of insecticide development turning into use of construction set composed of nitrogenous bases and genomic sequences of insect pests as instructions for assembling effective and safe oligonucleotide insecticides against sternorrhynchans and other groups of pests. Oligonucleotide insecticides are fulfilling a 90-year-old dream of creation of potent and selective chemical insecticides with highly adaptable structure to microevolution of insects.

## Author Contributions

Conceptualization, V.O.; methodology, V.O., N.G. and E.Y.; software, N.G.; validation, V.O., N.G., A.S. and Y.P.; formal analysis, V.O. and N.G.; investigation, V.O., N.G. and E.Y.; resources, V.O. and A.S; data curation, V.O. and N.G.; writing—original draft preparation, V.O., N.G., I.N., A.S., E.Y., E.L. and Y.P.; writing—review and editing, V.O., N.G., I.N., A.S., E.Y., E.L. and Y.P; visualization, V.O. and N.G.; supervision, V.O. and N.G.; project administration, V.O. and N.G.; funding acquisition, V.O. All authors have read and agreed to the published version of the manuscript.

## Institutional Review Board Statement

Not applicable.

### Informed Consent Statement

Not applicable.

## Data Availability Statement

The data presented in this study are openly available in [repository FigShare] at [10.6084/m9.figshare.28938248].

## Acknowledgments

We thank our many colleagues, too numerous to name, for the technical advances and lively discussions that prompted us to write this brief research report. We apologize to the many colleagues whose work has not been cited. Experiments were carried out at the Molecular Genetics and Biotechnologies Lab created within the framework of a state assignment V.I. Vernadsky Crimean Federal University for 2024 and the planning period of 2024–2026 No. FZEG-2024– 0001. We are very much indebted to all reviewers and our colleagues from the lab on DNA technologies, PCR analysis, and creation of DNA insecticides (V.I. Vernadsky Crimean Federal University, Department of General Biology and Genetics), and OLINSCIDE BIOTECH LLC for valuable comments on our manuscript.

**Disclaimer/Publisher’s Note:** The statements, opinions and data contained in all publications are solely those of the individual author(s) and contributor(s) and not of MDPI and/or the editor(s). MDPI and/or the editor(s) disclaim responsibility for any injury to people or property resulting from any ideas, methods, instructions or products referred to in the content.

